# Deciphering the rules of mRNA structure differentiation in vivo and in vitro with deep neural networks in Saccharomyces cerevisiae

**DOI:** 10.1101/337733

**Authors:** Haopeng Yu, Wenjing Meng, Yuanhui Mao, Yi Zhang, Qing Sun, Shiheng Tao

## Abstract

The structure of mRNA *in vivo* is influenced by various factors involved in the translation process, resulting in significant differentiation of mRNA structure from that *in vitro*. Because multiple factors cause the differentiation of *in vivo* and *in vitro* mRNA structures, it was difficult to perform a more accurate analysis of mRNA structures in previous studies. In this study, we have proposed a novel application of a deep neural network (DNN) model to predict the structural stability of mRNA *in vivo* by fitting six quantifiable features that may affect mRNA folding: ribosome density, minimum folding free energy, GC content, mRNA abundance, ribosomal initial density and position of mRNA structure. Simulated mutations of the mRNA structure were designed and then fed into the trained DNN model to compute their structural stability. We found unique effects of these six features on mRNA structural stability *in vivo*. Strikingly, the ribosome density of the structural region is the most important factor affecting the structural stability of mRNA *in vivo*, and the strength of the mRNA structure *in vitro* should have a relatively small effect on its structural stability *in vivo*. The recruitment of DNNs provides a new paradigm to decipher the differentiation of mRNA structure *in vivo* and *in vitro*. This improved knowledge on the mechanisms of factors influencing mRNA structural stability will facilitate the design and functional analysis of mRNA structure *in vivo*.

## Introduction

mRNA is a key component of the translation system. It not only carries protein coding information but also serves as an important vector for translation regulation information by folding into mRNA structures[1]. Notably, mRNA structures at the 5’UTR and ribosome binding site (RBS) regions have a significant effect on translation efficiency[2, 3]. Recent studies have shown that RNA structures *in vivo* are quite different from structures *in vitro* (as revealed by DMS probing[4, 5], DMS-MaPseq[6], icSHAPE[7] and SHAPE-MaP[2]). Some general conclusions have been drawn: the number of *in vivo* mRNA structures is far smaller than that *in vitro*[4], and mRNA structures in highly expressed genes, especially long-range base pairing structures, appear to be destabilized by translation[2]. However, the further investigation of these differences in the specific and accurate analysis of each mRNA structure is challenging, owing to the dynamic and unique nature of the mRNA structure itself; that is, each mRNA structure has a distinctive individual structure.

For translation by the ribosome to proceed smoothly, the mRNA structure must first be unwound, and a complex interaction between the translating ribosome and the mRNA structure occurs during translation. Translating ribosomes are one of the most important factors that cause structural differences *in vitro* and *in vivo*. Ribosome profiling is a method for determining the exact position of ribosomes in the transcriptome during translation with single-nucleotide resolution by deeply sequencing ribosome-protected mRNA fragments[8, 9]. This approach can effectively resolve the interactions between translating ribosomes and mRNA structure during translation. For example, the mRNA structure in the CDS region influences cotranslational protein folding by affecting the efficiency of ribosome translation or, in more extreme cases, by causing ribosomal pauses[10, 11, 12]. Additional factors that affect the mRNA structural stability *in vivo* could be primarily classified into several categories: gene features, such as mRNA abundance and translation initiation rate; structural features, such as the minimum free energy of structural subsequences; and other factors, such as the location of mRNA structure within a gene. Due to the multifaceted influence of these factors, the precise prediction of mRNA structural stability is a great challenge.

Deep learning, or artificial neural networks, is a type of machine learning that solves complex problems by learning “big data”. The deep neural network (DNN), a type of deep learning, has multiple hidden layers and units that can model complex nonlinear relationships and is widely used in natural language processing[13], speech recognition[14] and the sensational AlphaGo[15]. Deep learning is currently being applied to biological “big data”, such as the detection of breast cancer based on histological images[16], the analysis of ribosomal stalling sites based on ribosome profiling[17] and the prediction of DNA- and RNA-binding proteins[18]. We have proposed a novel application of the DNN model to fit mRNA structure features and subsequently predict the structural state *in vivo*, revealing the impact of these features on the *in vivo* mRNA structural stability.

As the simplest eukaryote, *S. cerevisiae* was first and most commonly used in research on *in vivo* mRNA structure and ribosome occupancy[4, 5, 8]. Therefore, in this study, *S. cerevisiae* was selected as the research species for a series of studies on the analysis of mRNA structure *in vivo*. Among experimental approaches to detecting RNA structure *in vivo*, DMS probing was demonstrated to be effective for the determination of mRNA structure in *S. cerevisiae* by setting threshold values of the Gini coefficient and Pearson correlation[4]. Thus, we proposed identifying *in vitro* and *in vivo* mRNA structures by this method and dividing them into two classes (0 and 1, structures with stable and unstable tendencies *in vivo*). Then, the DNN model was trained with these data. After obtaining a precise model to predict the structural state *in vivo*, we can mutate different features of these mRNA structures and decipher the specific impact of these features on mRNA structural stability (Fig 1).

**Figure 1.**
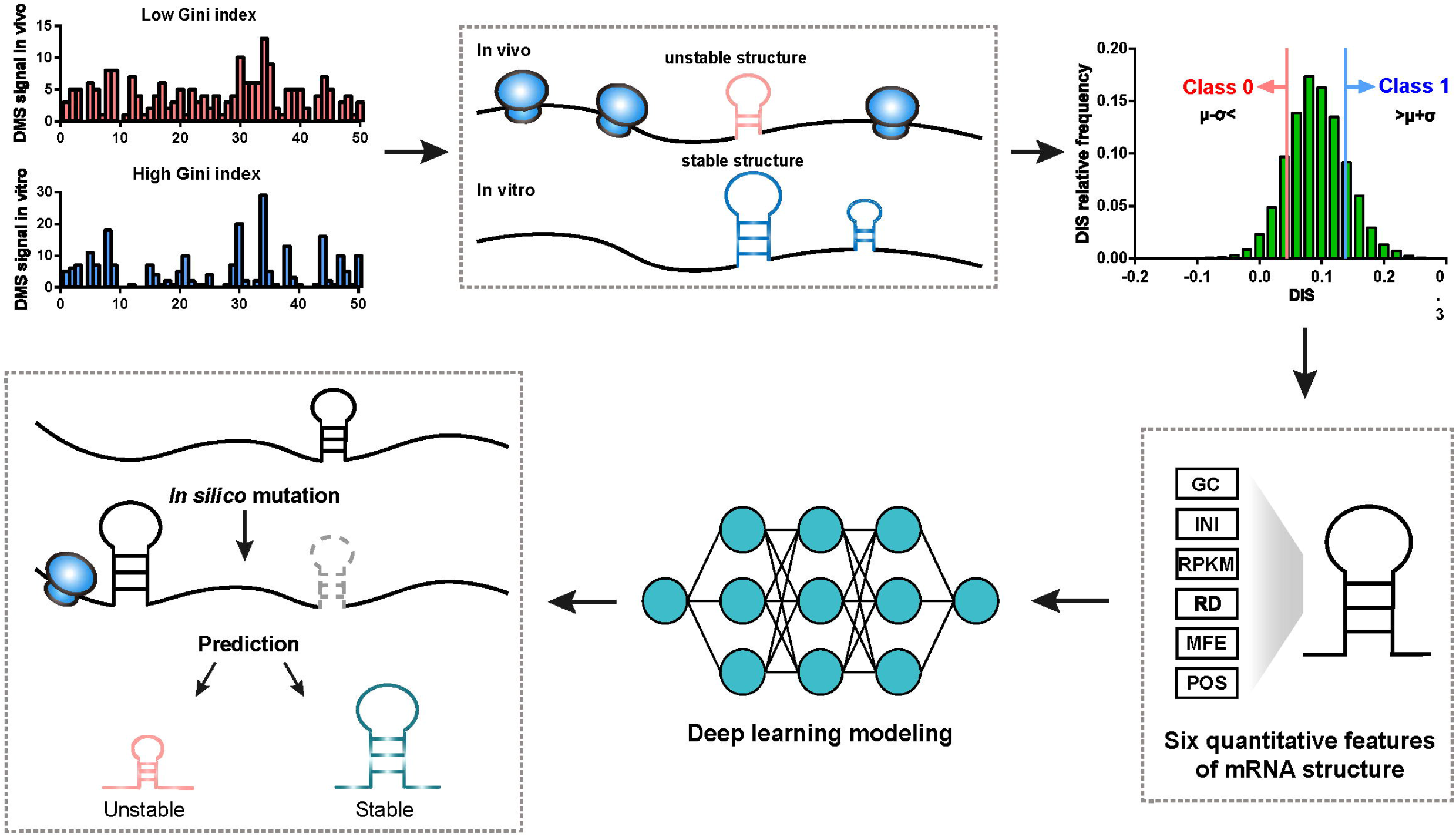
Schematic overview of deep learning modeling and prediction pipeline. The Gini index was calculated to measure the stability of the mRNA structure. The difference between the Gini index values *in vitro* and *in vivo* was used to classify the mRNA structures into class 0, manifesting stable tendencies, and class 1, manifesting unstable tendencies. Then, the two classes were fed into the DNN model for training. After the trained model was obtained, various mutations of mRNA structures were designed, and the model was used to predict whether they would be stable.

## Results

### Constructing a deep learning model with 6 mRNA structure features

We proposed a DNN model that fits mRNA structure differentiation *in vitro* and *in vivo* with quantifiable features that may influence structural stability, thereby realizing the ability to predict mRNA structural stability *in vivo* (Fig 1). Unlike previous mRNA structure prediction models that aimed at constructing RNA structures[19], this deep learning model was mainly used to predict whether mRNA structures *in vivo* were stable or unstable. DMS profiling[4] is a chemical probing method that identifies methylated adenine and cytosine bases in single-stranded RNA and can be used to provide a global feature describing *in vivo* and *in vitro* structures as well as denatured mRNA structures by measuring the Gini index[20] and Pearson correlation coefficient. A higher Gini index corresponded to a more stable structural region, and a lower index corresponded to a less stable structural region. Normally, due to a variety of factors, such as translating ribosomes, the *in vitro* structure was usually more stable than that *in vivo*[4]. Therefore, the ‘disappearance trend’ (DIS), which was equal to the Gini index of mRNA structures *in vitro* minus that *in vivo*, was used to measure the differences in mRNA structure *in vivo* and *in vitro*. In this study, the *in vitro* mRNA structures of *Saccharomyces cerevisiae* were all divided into two classes by DIS. Class 1 contained structures beyond the mean plus standard deviation (SD), which represented structures in unstable states. Class 0 was structures below the mean minus SD, which represented structures in stable states (Fig 1). After multiple thresholds of classification were tested, one SD was determined to ensure the appropriate size of training sets and deep learning model accuracy.

Stable and unstable mRNA structures could appear on the same gene, and several differences in structural features might be interpreted as corresponding to the existence of these two states (Fig 2a). Previous studies found that a variety of factors can affect mRNA structure *in vivo*, including ribosome density (RD), gene expression, tRNA levels and the structural stability of the mRNA itself[2, 11, 12, 21]. Six quantitative features were chosen for modeling: gene features, including reads per kilo base per million mapped reads (RPKM) of RNA-Seq, which implied the mRNA abundance, and the initial translating ribosome density (INI), calculated from ribosome profiling data; subsequence structure features, including the minimum free energy (MFE) and GC content (GC) of the subsequence, which reflect mRNA structure stability; and other features, including the relative position of the structure in the gene (POS) and the ribosome density of the structure region (RD). Notably, the RD and INI values in this project were averaged from five different studies of ribosome profiling data in wild-type yeast[22, 23, 24, 25, 26]. The DIS value of mRNA structure showed markedly positive relationships to RD and RPKM, implying that high ribosomal density and high mRNA abundance were not conducive to the formation of stable mRNA structure (Fig 2b). These results were consistent with observations from prior studies[2, 27]. The six features all showed statistically significant differences between the two mRNA structure classes. However, the difference was visible only in the RD, RPKM, and GC parts (Fig 2c). It could be initially concluded that these six features do not individually have strong effects on the mRNA structural stability, and it was difficult to resolve their contributions to mRNA structural stability from basic statistics. Therefore, we adopted DNN modeling to decipher the relationships between these complex regulatory factors.

**Figure 2.**
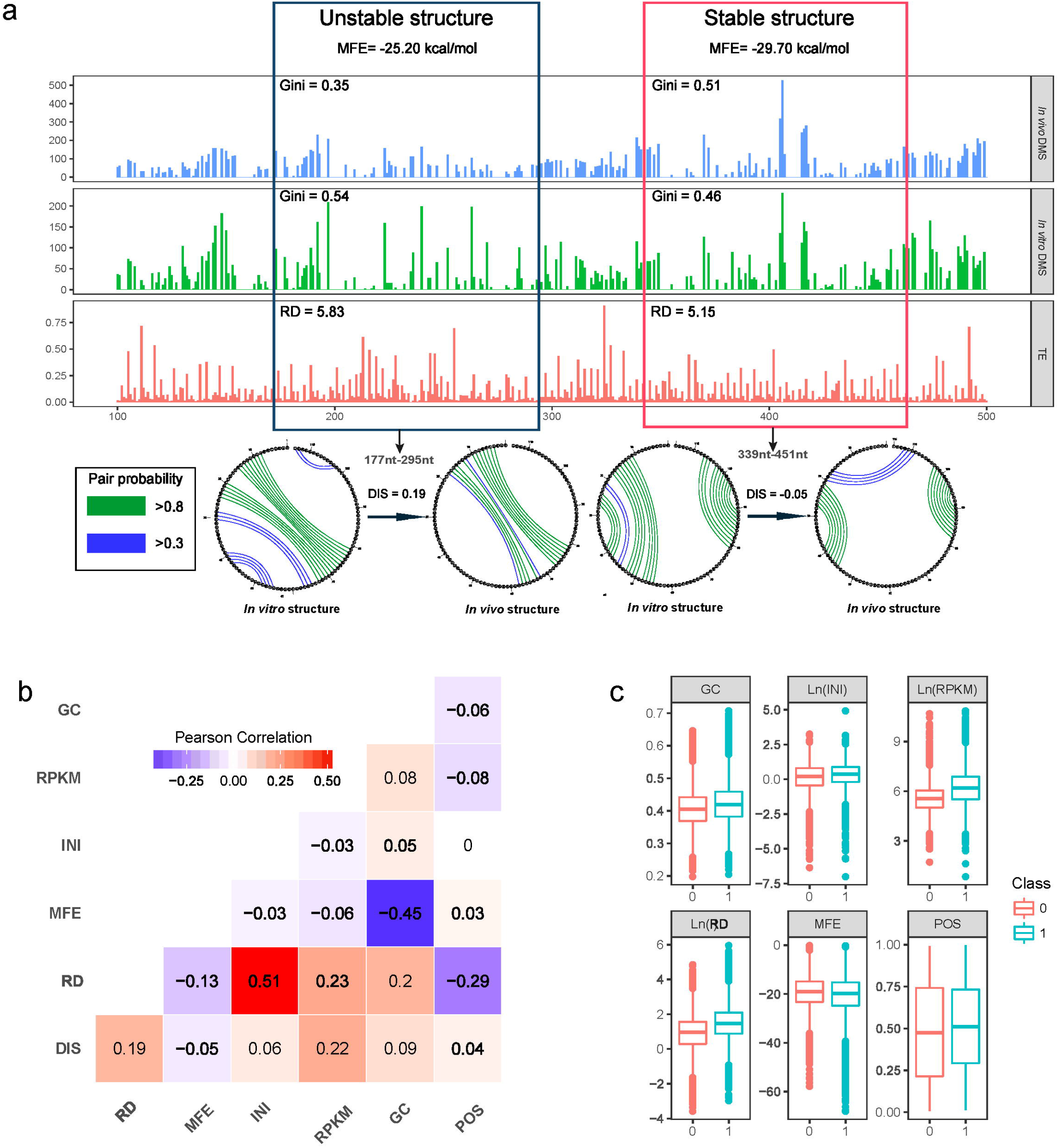
Stable and unstable mRNA structures with various structural features. (a) Stable and unstable structures with different structural features in the YML110C gene (Unstable structure: 117nt-295nt, stable structure: 341nt-455nt). The circular mRNA structure map shows the difference in in vitro and in vivo structures of stable and unstable structures. (b) The Pearson correlation coefficients between the six characteristic structural features. (c) Distribution of mRNA structural features between the two classes.

We established the DNN sequential model using the TensorFlow deep learning framework[28], named DeepRSS (using a **Deep** learning approach to predict m**R**NA **S**tructural **S**tability *in vivo*.). By performing a 10-fold cross-validation on a variety of hyperparameters, the final end-to-end DNN model has 9 fully connected layers with 256 cells per layer; the activation functions adopted were ReLU [29] and Sigmoid; the Adam optimization function[30] was adopted to accelerate the training process; and optimization techniques, batch normalization[31] and early stopping[32] was added to prevent overfitting of the model (Fig 7, Table S1). The precision of the DeepRSS model reached 99.71% and area under ROC curve (AUC) reached 0.998001. Thus, a DNN model that can accurately predict the *in vivo* mRNA structure state from *in vitro* structural features was established.

### High ribosome density resulted in instability of the mRNA structure

Translating ribosomes are a critical factor that affects mRNA structural stability. Previous studies have demonstrated a negative correlation between ribosomal translation efficiency and in vivo mRNA structure stability; that is, high translation efficiency corresponds to low structural stability in a region ^2,5,27,32^. We performed single-factor 100 times gradient mutations on the ribosome density (RD) of the whole mRNA structures, containing 68,929 unstable structures and 66,915 stable structures. These gradient mutations actually simulated the *in vivo* structural state of an mRNA structure under different RD conditions. By predicting the state of the mutated mRNA structures using DeepRSS, a clear correlation between the RD and the average structural transformation ratio was shown, that is, high RD caused stable mRNA structures to become unstable, while low RD could stabilize unstable structures (Fig 3a), which supported the previous study. This correlation is most pronounced for intervals greater than −2.0 (corresponding to 0.48 before normalization) and less than 3.0 (corresponding to 4.07 before normalization). Due to the steric hindrance of the translating ribosome, if the RD of the mRNA structural region was high, the ribosome would occupy the folding space of the structural region, thus causing the mRNA structure to not be formed.

**Figure 3.**
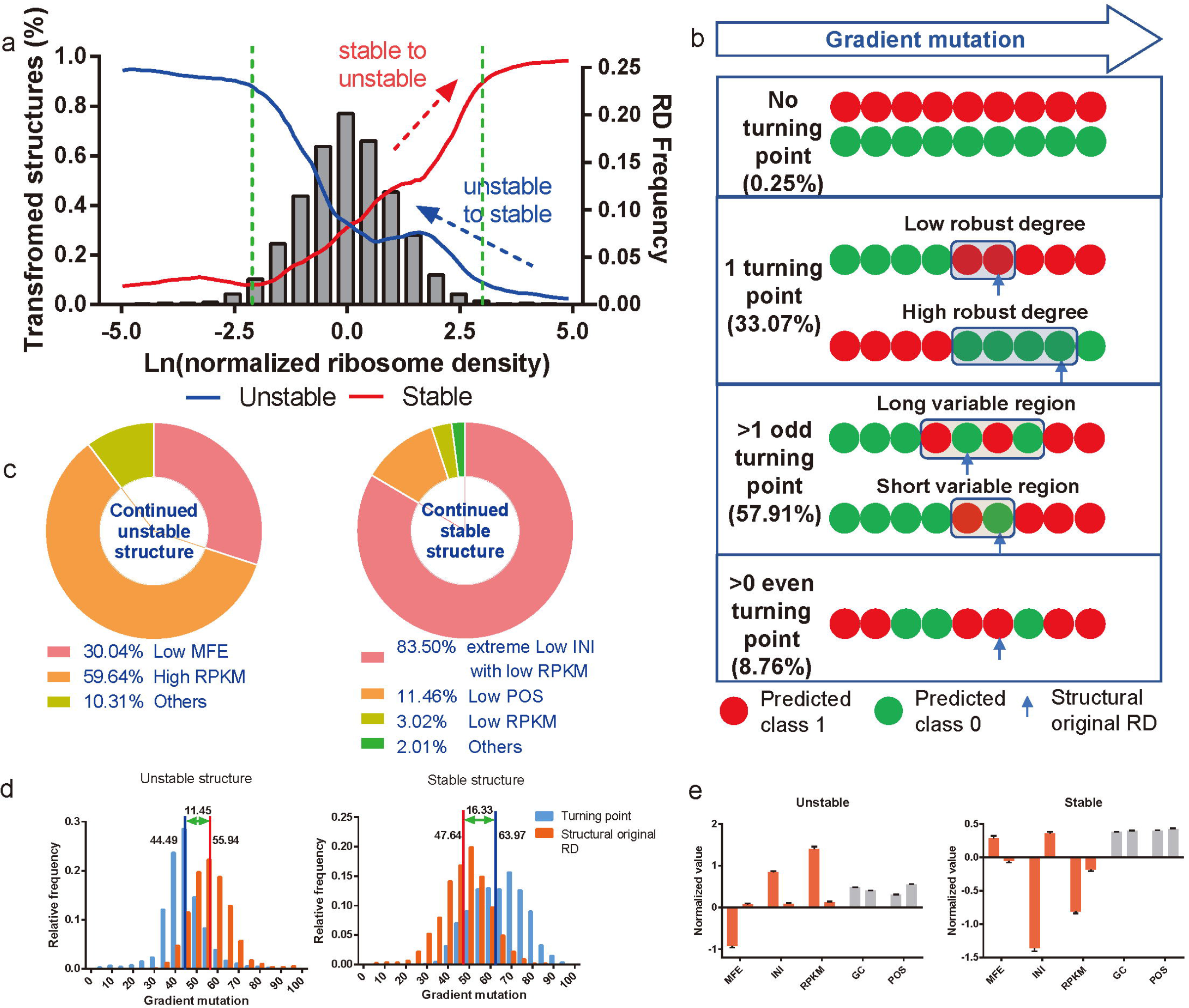
The effect of ribosome density on the structural state of mRNA in vivo. (a) Single-factor mutation of ribosome density. The ordinate was the ratio of transformed structural state. When the value of transformed structures reached 1, it represented that the all structures transformed to the opposite of their original state. Structural state transformed from unstable to stable, named “Unstable to stable”, and vice versa named “Stable to unstable”. THe histogram of the RD frequency distribution was also shown. (b) Different turning points in gradient mutations of RD. (c) Venn diagrams that characterized the main 4 structural features of continued stable and unstable structures. (d) The distribution of the turning point and the structural original RD position in gradient mutations. The blue and red lines represented the mean of the turning point position and the structural original RD position.

The *in vivo* mRNA structures were classified according to the predicted results of the single factor gradient mutations of RD (Fig 3b). In the gradient mutation process from small to large, if the mRNA structure state changed, it was considered a turning point. If the mRNA structural state did not change in the gradient mutation of RD, i.e., there was no turning point in the result of 100 mutations, then this mRNA structure was characterized by ‘continued stable’ or ‘continued unstable’ structures. The continued unstable structures were located on genes with high mRNA abundance and, more interestingly, these structures showed high GC content and low MFE, that is, they were a stable structure *in vitro* (Fig 3c). It can also be seen that continued unstable mRNA structures originally had a high ribosome density. On the other hand, the continued stable structures were located on genes with low translational initiation ribosome density or low mRNA abundance, as opposed to continued unstable structures. The features of *in vitro* mRNA structures, including the GC content and MFE, are described in detail in the subsequent results. In terms of the translation level of the gene itself, including RPKM and INI, mRNAs with high mRNA abundance often need to maintain high protein expression levels, and the mRNA structure as an energy barrier hinders ribosome translation; thus, they are difficult to stabilize. However, genes with low INI and low RPKM do not have a heavy ‘translation task’, and can be folded and persist in a stable state.

mRNA structures with one turning point accounted for 33.07% of the total mRNA structures. We calculated the distance between the position of the original structural RD at 100 mutations and the position of the turning point as an evaluation of whether the structural state was easily interfered by the RD variation, termed the ‘robust degree’ (Fig 3b). In general, the unstable structure was on average higher than its turning point of 11.45, while the stable structure was on average lower than its turning point of 16.33 (Fig 3d). That is, an unstable structure was more susceptible to the structural state due to the influence of RD fluctuations than a stable structure. After sorting according to the robust degree, the mRNA structures of the top 1,000 (structures of high robust degree) and the last 1,000 (structures of low robust degree) were extracted. Comparing the high and low robust degree structures, a more stable *in vitro* structure (lower MFE), higher initial ribosome density (higher INI), and higher mRNA abundance (higher RPKM) allowed unstable structures to maintain their unstable state. Conversely, an unstable *in vitro* structure, lower initial ribosome density, and lower mRNA abundance allowed stable structures to maintain their state (Fig 3e). The mRNA structures with odd turning points greater than 1 accounted for 57.91% of the total structures, and slightly different from only one turning point, their structural state fluctuated within a certain RD interval, named a ‘variable region’. The regularity of these mRNA structures for RD fluctuations was the same as that of only one turning point structures, that is, the characteristics of the gene and its own MFE determine the length of these ‘variable regions’.

### mRNA structure prone to stability at the 5’-end of the gene

Our trained model provided accurate predictions of the yeast mRNA structural state *in vivo* through six structure features. Here, we sought to establish the relationship between mRNA structure and its relative position in the gene CDS region. Previous studies analyzed the *in vitro* mRNA structure of various species to derive a tendency for stable RNA structures to form downstream of the translation initiation codon (AUG), which might favor ribosome formation in the ‘ramp’[33, 34, 35]. This feature could also be seen from the statistics of the 2 classes (stable and unstable) of mRNA structure in yeast, with a higher proportion of stable mRNA structures at the 5’ and 3’ ends of the gene (Fig S1). According to the ratios of different mRNA abundance genes of two classes (class 0 and class 1), it was found that high mRNA abundance was more likely to cause structural instability, while there was small difference for low mRNA abundance (Fig 4b). We changed the relative position (POS) of the mRNA structure on the gene by gradient mutations and found that the state of 85.00% originally unstable structures transformed to stable after transferring structures downstream of the AUG (Fig 4a). As the mRNA structure was transferred to the upstream gene, the proportion of stable structures increased. The mRNA structure of the middle part of the gene remained basically unchanged, and the stable state of the structure at the 3’ end (beyond 0.8) revealed a slight trend toward stable transformation, especially to unstable structures. The mRNA structure tended to form *in vivo* at the 5’ and 3’ ends of the gene, which was consistent with the findings of previous studies[33, 35]. The proportion of no turning point structures in the POS gradient mutations was much higher than that in the RP gradient mutations, with 6.27% unstable structures and 16.57% stable structures. Classified from the point of the turning point, the structural region was occupied by a higher RD, and the structure located in the highly expressed gene was also a cause of the continued unstable structure (Fig 6a). Located in low expression genes, the absence of ribosomal coverage also contributed to the continued stable structures by positional mutations. Furthermore, a proposed POS and RD co-mutation was designed by mutating these two features as the first structure of the 5’ end of the gene, and a similar result was obtained (Fig 4c).

**Figure 4.**
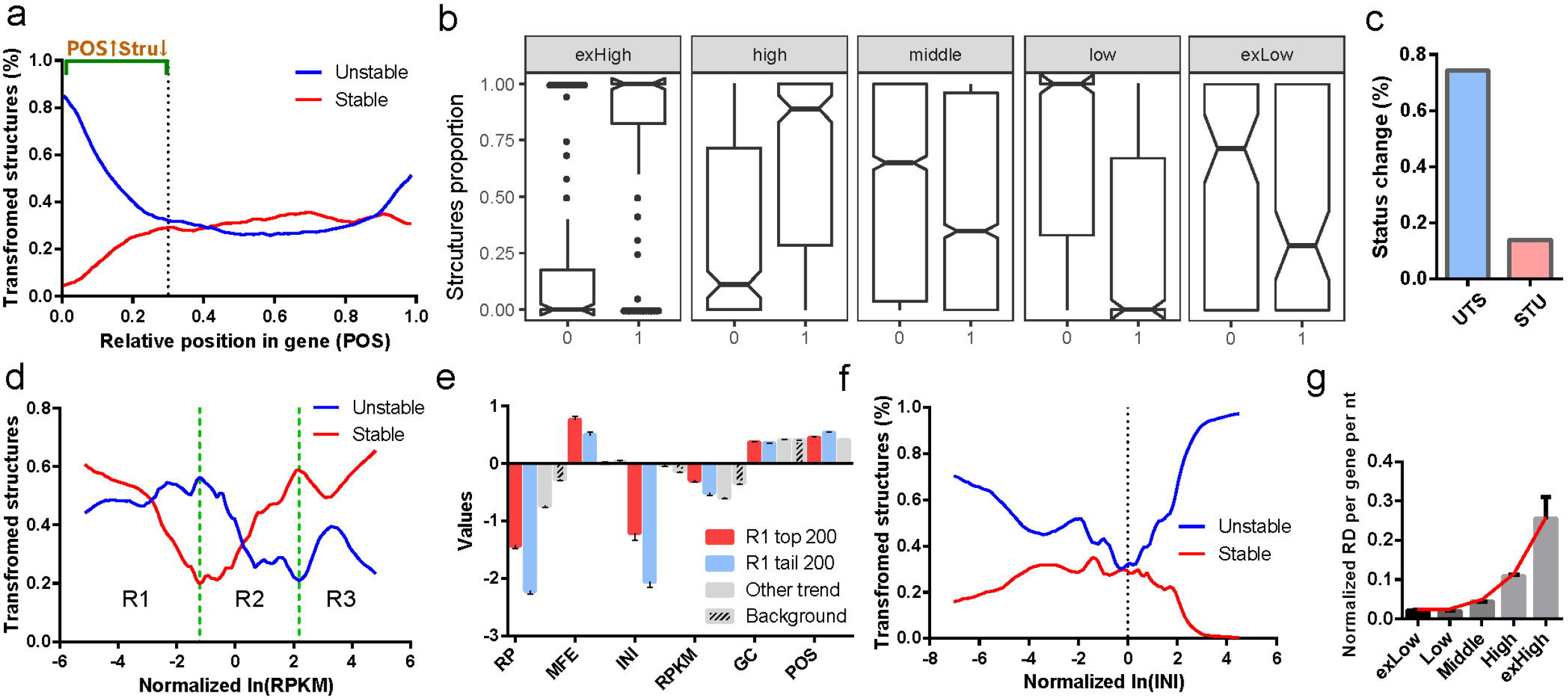
The effect of gene characteristics on mRNA structural stability. (a, d, f) Single-factor gradient mutation of POS, RPKM and INI in mRNA structure (b) The ratio of stable and unstable structures on a gene for different groups of RPKM. (c) Transformation ratio of unstable to stable mRNA structures (UTS) and stable to unstable structure (STU) after co-mutation of the mRNA structural position and RD. (e) Structural features of the one turning point mRNA structure with R1 trend of RPKM. “Other trend” is another structure with one turning point. (g) Normalized RD per gene per nt of genes of different INI groups.

### Effects of mRNA abundance and initiation RD on mRNA structure stability

Two primary characteristics of the gene affect the structural stability: mRNA abundance (RPKM value in this project) and translation initiation ribosome density (INI value in this project, which could be translated into the translation initiation rate through data processing). The features, RPKM and INI, were obtained from RNA-Seq data and ribosome profiling data, respectively. The mRNA abundance represented the mRNA copy number, which has a certain correlation but is not equal to the protein synthesis efficiency obtained by ribosome profiling[36, 37, 38]. According to our understanding, the mRNA structure serves as a barrier to ribosome translation and remaining stable is not necessarily compatible with the translation efficiency of highly expressed genes. As there was no strong “translation task” for low-expression genes, structural instability was not strongly needed; hence, these structures tended to be stable. However, in the process of single-factor mutations in the mRNA structure of RPKM, an interesting “V” trend was found, as shown in Fig 4d. Based on the trend of the mRNA structural transformation, we divided the RPKM values into three intervals, R1 to R3. Among them, the trend of region R2 was the same as we speculated, that is, as the number of unstable structures increased with the increase of the RPKM value, the number of stable structures decreased. The most interesting structural transformation region was R1. When the RPKM dropped to a certain extent (normalized ln(RPKM)<-1.21), many of the unstable structures had a hard time being converted into stable structures, but the originally stable mRNA structure became unstable due to this decrease. In order to explain this trend in the stable structures, only one turning point structure in which the turning point appeared in the R1 area was selected, and the top 200 and last 200 structural features were compared after turning point position sorting. The “Top 200” are the 200 structures closest to the V-shaped turning point, representing the structures that are most likely to form the R1 trend, and the “Last 200” are the opposite. The mRNA structures with the R1 trend mainly had a lower RD, a more unstable *in vitro* structure and a lower initial ribosome density than the other trend and background groups (Fig 4e). Among them, the “Top 200” structures with the R1 trend had a higher RD, a more unstable external structure and a higher INI than the “Last 200”. It should be noted that in the process of decreasing the RPKM mutation, the RD was increased due to the decrease in transcription copies. As we discussed earlier, the structure becomes unstable due to an increase in the RD, thus causing a trend in R1. The “Top 200” structures exhibited this trend earlier than the “Last 200” structures because of their higher RD and more unstable *in vitro* structures. Subsequent calculations found that fluctuations in the R3 region were also associated with decreased RD due to increased RPKM.

Translation initiation is the rate-limiting step in the translation process and is regulated by a variety of factors including the mRNA structure. The mRNA structural feature, INI, which is the sum of the normalized ribosome densities relative to the start codon 0 to +15 region, can reflect the translation initiation efficiency. INI single-factor gradient mutations showed that INI tended to stabilize the structure above and below the mean, and this “V” trend was more obvious for the originally unstable structure (Fig 4f). It could also be seen that the ribosome density per gene per nt and INI were positively correlated in these five groups (Fig 4g). That is, when the INI was decreased, the RD was also decreased, thus making the mRNA structure more stable. However, increased INI also leads to an increase in RD, but the mRNA structure is more stable. It is possible that because of the reduced efficiency of translation initiation, the ribosome interval on the mRNA increases; thus, the local mRNA structure has time to re-form the folded re-formation structure. This hypothesis has also been proposed in previous studies by computer simulation of the ribosome translation process [21]. We also tried to mutate two features at the same time, mRNA abundance and translation initiation rate, but the structural transformation trend fluctuated greatly, and no obvious regular behavior could be identified.

### Effect of the mRNA structure subsequence composition on its stability

The minimum free energy (MFE) and GC content were used to evaluate the stability of the mRNA structure *in vitro* based on its sequence composition[21, 33, 39, 40]. MFE was significantly negatively correlated with GC, and the Pearson correlation reached −0.45 (Fig 2b). Compared to mutations in the other features discussed previously, the results of single-factor mutations in both MFE and GC had very distinctive trend (Fig 5a, 5b). The transformed mRNA structure curve showed a V-shaped trend with a mean value as the demarcation point. As the changes in the normalized MFE value moved in both directions from the mean 0, both stable and unstable structures would shift toward the opposite state. At the MFE and GC mean points, the lowest point of the structural transformed rate, that is, the mRNA structure at the mean, can more easily maintain its original state. This trend actually reflects the fact that the *in vitro* mRNA structure has a limited effect on the mRNA structure *in vivo* because this trend was similar to its distribution. It also can be seen from Fig 5a and Fig 5b that even extreme mutations to the MFE and GC content cannot change the mRNA structural state by more than half. Because there are many factors in the *in vivo* environment that can affect structural stability, we believe that the sequence composition is not the most important factor in determining the stability of this structure *in vivo*. The MFE and GC content are interrelated and related to the sequence composition; therefore, mutations designed to target both sequences need to be based on specific sequences. To reduce the impact of other structural factors, we selected two mRNA structures from all the structures, of which the six structural features were close to the mean. Random mutations of the structural sequence were made and 16 mutations in which the absolute value of the MFE and GC were simultaneously increased or decreased were selected. It can be seen that if the effects of other structural features were attenuated, the direction of the mRNA structural change was directly affected by the MFE and GC (Fig 5c). An increase in the MFE and a decrease in the GC content correspond to a decrease in the number of mRNA structure pairs and the pairing probability, which corresponds to the tendency of the mRNA to be structurally transformed (Fig 5d). When the interference of other factors is removed, the sequence composition of the mRNA structure directly affects the stability of its structure *in vivo*.

**Figure 5.**
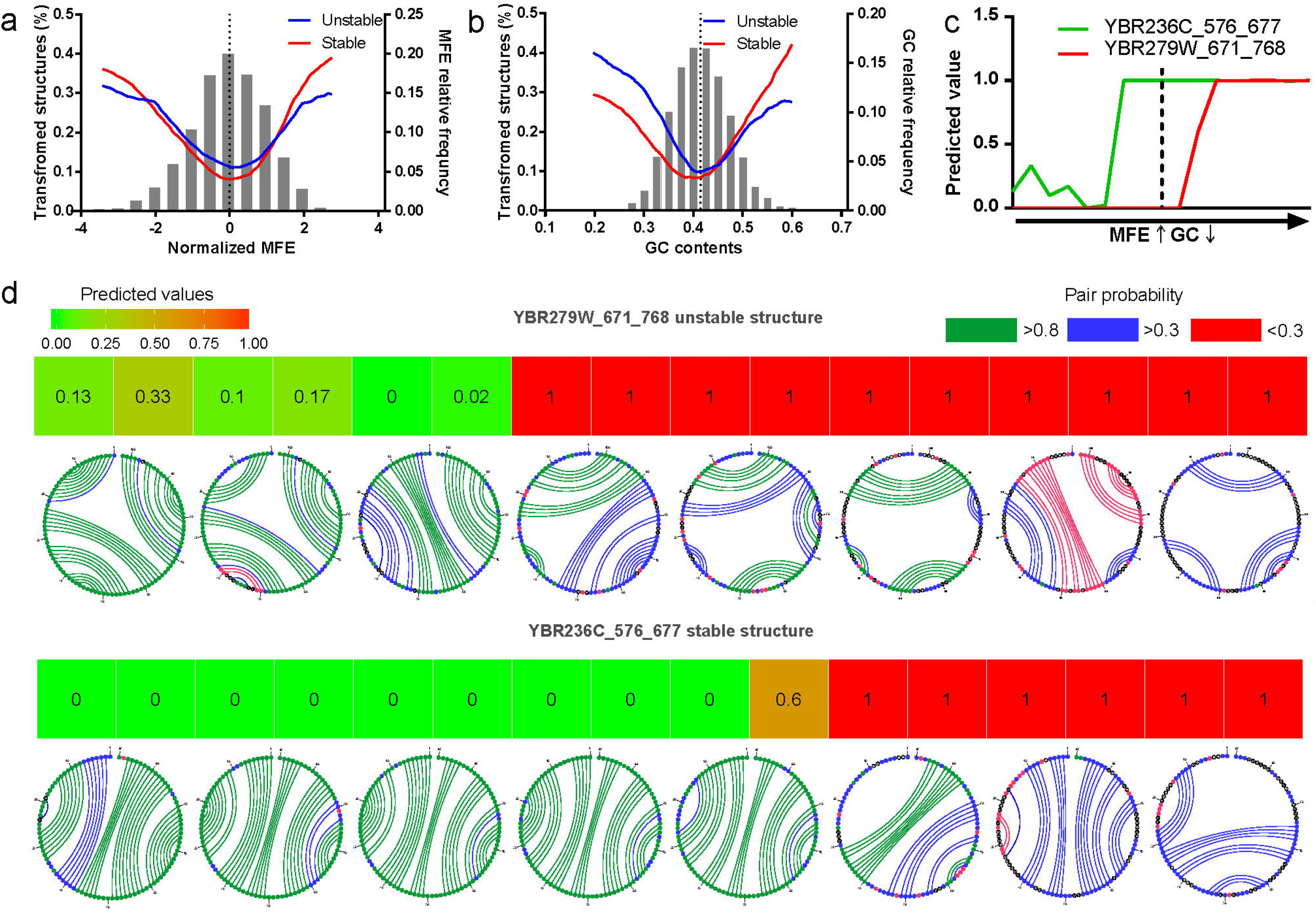
The effect of MFE and GC content on structural stability in vivo. (a, b) Single-factor mutation of MFE and GC. (c) The effect of 2 mRNA structures on the co-mutation of MFE and GC. Green is the originally unstable structure, red is the original stable structure, and the dotted line is the original MFE and GC of these two structures. (d) Detailed process of two mRNA structural mutations and structural state transitions. The predicted values are predicted by the DeepRSS model, <0.5 can be classified as a stable structure, and >0.5 is classified as an unstable structure. In vitro mRNA structural pairing different mutational stages is shown, with different colors representing different pairing probability.

### The stability of the mRNA structure in vivo is mainly related to intracellular factors

The structural ribosomal density (RD), the structural minimum free energy (MFE), the ribosome density (INI) in the translation initiation region, the mRNA abundance (RPKM), the GC content of the structural region sequence (GC) and the relative position of the mRNA structure in the gene (POS) are six important factors affecting the structural stability of the body. In terms of the 6 structural features in gradient mutations, the reason for continuously stable or unstable structures was determined by setting the standard deviation of the structural feature to a threshold value (Fig 6a). Overall, the causes of structural instability are mainly high RD, high RPKM and high INI. Stable *in vitro* structures (high GC and low MFE) also appear to be responsible for their instability, but according to the previously mentioned conclusions, this phenomenon may have been determined to be unstable due to other factors (such as high RP or high RPKM) and is therefore not selected during evolution. The factors that contribute to the stability of the mRNA structure *in vivo* are mainly the low RD of the structural region, low RPKM, low INI and a structural position closer to the 3’ end of the 5’ end. To measure the extent to which the six structural features affect the mRNA structure, we defined the “impact ratio”, which is based on the proportion of continued stable or unstable structures in the gradient mutation. It can be concluded that the factors affecting the structural stability *in vivo* are, from highest to lowest, the RD, RPKM, INI and POS. That is, these factors are more likely to cause the mRNA structure to change its original state. Among them, INI and POS have stronger effects on unstable structures than on stable structures. MFE and GC have the lowest effect on the stability of the mRNA structure *in vivo*. In addition, we calculated the tolerance of phylogenetically conserved mRNA structures to these structural mutations. Phylogenetically conserved mRNA structures were evidenced by DMS-seq and inspected by phylogenetic conservation analysis[4]. Positive and negative mutations in three features of mRNA structure, RD, RPKM and INI, revealed that the conserved mRNA structure was better able than random structures to maintain its own state of stability when subjected to the same degree of mutation (Fig 6c). Consequently, the conserved structure was more tolerant to fluctuations in the intracellular environment than the random structure.

**Figure 6.**
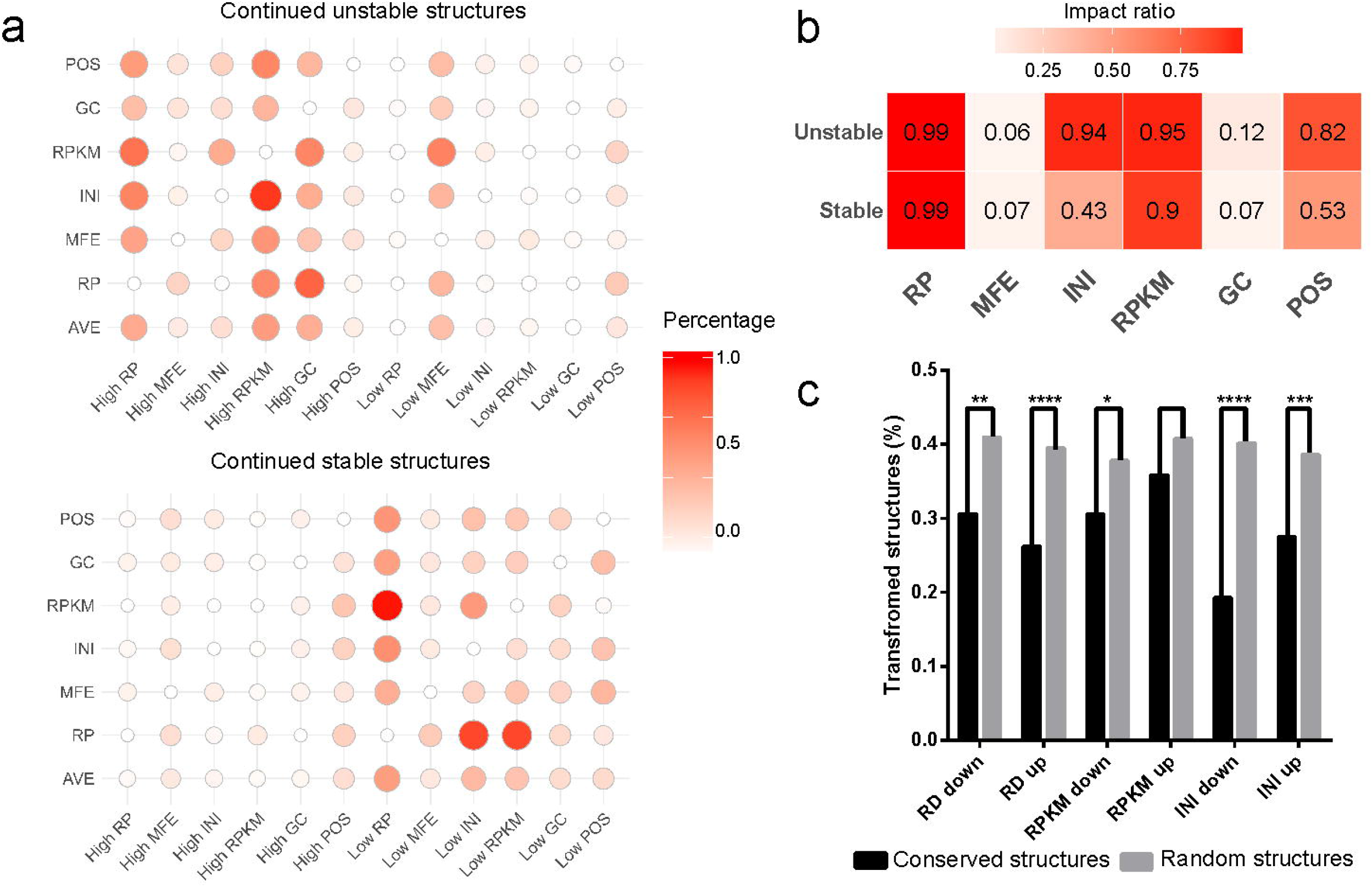
The effect of six structural features on structural stability in vivo. (a) The reason for the continued stable and unstable structure is produced in the gradient mutation of 6 structural features. (b) The impact ration of six structural features on stable and unstable mRNA structures. (c) Evolutionary conserved structure and random structure tolerance to mutations.

## Discussion

Since the invention of large-scale, high-throughput methods for *in vivo* mRNA structure determination, previous studies have statistically derived general conclusions regarding *in vivo* and *in vitro* mRNA structural differences, such as that the number of *in vitro* mRNA structures was greater than that *in vivo*, and the structural differences *in vivo* and *in vitro* mRNA were mainly caused by ribosome translation[2, 4, 5, 6, 7]. Because multiple factors caused differences in *in vivo* and *in vitro* structures, it was difficult to perform a further accurate or “personalized analysis” of mRNA structures in previous studies.

In this study, we used a novel deep learning application to accurately model labeled mRNA structures in yeast and six quantitative structural features (RD of structure regions, MFE and GC content of subsequences, gene mRNA abundance, ribosomal initiation density, and relative structural position). Through extensive model selection and optimization, we obtained the DeepRSS model, which best predicted the structural state *in vivo*. The second classification prediction accuracy of the trained DeepRSS model reached 99.71% and there was no over-fitting. The rules explaining the effects of these six features on mRNA structural stability were then resolved by performing *in silico* mutations in these six features and subsequent prediction by the DeepRSS model. We found that the RD reduces the stability of the mRNA structure in vivo, while unstable mRNA structures could also be converted to stable structures *in vivo* by reducing the RD. The RD of the structural region is the most important factor affecting the mRNA structural stability. The relative position of the mRNA structure on the gene also affects its stability; that is, the *in vivo* mRNA structure near the 5’ end (< 20%) in particular and the 3’ end (>80%) tends to be stable. Genes with high mRNA abundance require rapid translation to produce the desired protein product, and thus the mRNA structure of these mRNA or increasing the mRNA abundance *in silico* tends to cause mRNA instability. However, when the abundance of the mRNA is reduced to a certain extent by mutation, the structure may become unstable, which may be caused by a decrease in the density of ribosomes on the mRNA. The translation initiation efficiency of mRNA is an important factor affecting the translation efficiency of the entire mRNA, and it is positively correlated with the RD on the mRNA. Decreasing the RD in the translation initiation region tends to stabilize the mRNA structure, while a high RD of the translation initiation region increases the ribosome spacing on the mRNA and cause the mRNA structure to become stable. The sequence of the mRNA structural region determines the strength of its *in vitro* mRNA structure through the MFE algorithm or GC content, but has limited effects on the stability of its structure *in vivo*. Among the six structural features, high RD, high mRNA abundance and high translation initiation efficiency are the main causes of structural instability *in vivo*. The stable structure is mainly due to a low RD and low mRNA abundance, as well as its low initial RD and proximity to the 5’ end and 3’ end of the mRNA. Simultaneously, changing the structural state of phylogenetically conserved mRNA structures is more difficult than changing the structural state of random structures even if the same structural features are mutated.

This study is the first to use a deep learning approach to decipher the rules of mRNA structure differentiation *in vitro* and *in vivo*. DNNs were proven to be usable for solving complex regulatory issues in the translation process. The DNN model accurately predicted the relationships between these six mRNA structural features and the structural state and also resolved the range of influence of these features on structural stability. However, some limitations are worth noting. Previous studies have shown that codon bias and tRNA abundance could affect the ribosome translation rate and thus influence the structural stability of mRNA *in vivo*[12, 21, 41]. Codon bias and tRNA abundance were not considered separately in our model, because the ribosome translation information of these two factors was already carried by the RD data. Our results are encouraging and should be validated in a larger number of mutant yeast species. Future works should therefore focus on the following points. Codon biases and tRNA abundance, as separate features, should be used instead of RD to make the DNN model more universal and to perform an impact analysis of the mRNA structural stability. In addition, more species, such as Arabidopsis and mouse, as well as more detection methods, such as SHAPE-MaP and icSHAPE, should be considered in the state prediction of mRNA structure *in vivo*.

## Materials and Methods

### Raw data preprocessing

*Saccharomyces cerevisiae* RNA structure data were obtained from a DMS profiling experiment (containing *in vivo*, *in vitro* and denatured data sets) published in the NCBI GEO database, accession number GSE458034. We identified the DMS signals of the specified genome positions in each gene, trimmed G/T sites in the gene sequence, and then used sliding windows of 50 nt (full-length sliding windows, i.e., after the addition of G and T sites, were approximately 100 nt) to divide the trimmed genes into regions and filtered out regions with low DMS signal (<1 per nt). Gene sequences were retrieved from SGD database assembly R62. Multiple sets of ribosome profiling data (RP) were combined to approach the true *S. cerevisiae* ribosome occupancy. The RP data of wild-type yeast were based on the following studies: Zinshteyn and Gilbert *et al.* 2013[22], Lareau *et al*. 2014[23], Albert *et al*. 2014[24] (BY and RM samples), Young *et al*. 2016[25] and Nissley *et al*. 2016[26] (Rep1 and Rep2 samples). The RNA-Seq data were from Albert *et al*. 2014 (BY and RM samples)[24]. Then, the RP and RNA-Seq data were trimmed and mapped to the yeast genome (assembly R62) by Bowtie and normalized by the RPKM of the RNA-Seq data[42]. For the yeast fragments, the offset of 15 nucleotides from the 5’ end represented the P-site of the ribosome position[36]. Therefore, ribosome occupancy at nucleotide resolution was given by the location of the genome position of the 15th nucleotide of ribosomal fragments. The ribosome footprint of the corresponding position was obtained by summing all these RP data from different studies.

### Classification of mRNA structures by DMS probing data

We defined the RNA structure by two metrics: Pearson’s correlation coefficient (r<0.55) and the Gini coefficient (Gini<0.14), as described[4]. After removal of the G/T sites in the sequence, a 50 nt sliding window was used to divide the CDS regions of all genes. First, we determined the structure *in vitro* by thresholds of r and Gini. To more reasonably analyze the trend of disappearance of RNA structure, we proposed a novel measure for the mRNA structural regions, DIS, which was equivalent to subtracting the *in vitro* and *in vivo* Gini coefficients of the structural region (G). For the mRNA structural region of length *n*, the DMS signal per site is *S*_*i*_, and its DIS and Gini coefficient are as follows:

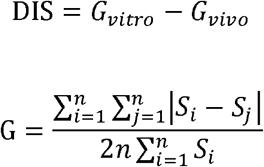

### Calculation and standardization of six features of mRNA structures

We chose six structural features as measures of mRNA structure. The feature TE (translation efficiency) was the protein synthesis rate of the structural region[43]; MFE was the minimum free energy of a local subsequence, calculated by the Vienna (v.2.1.9) package RNAfold[44] function and visualized by RNAstructure[45]; INI reflected the translation efficiency at the beginning of the gene region (+15 nt); RPKM represented the relative number of transcripts of a gene and was calculated from the RNA-Seq data; GC represented the GC content of the sequence of the structural region; and POS was the relative position of the structural region in the gene, which was obtained by dividing the sequence length by the central position of the structural region. Three of these features, TE, INI and RPKM, were exponentially distributed, so a logarithmic process was performed. Prior to deep learning training, the parameters (except POS and GC) were normalized ((x-μ)/σ^2^) to make the model more accurate.

### Established deep neural network model

We established a DNN model named DeepRSS (using a **deep** learning approach to predict the m**R**NA **s**tructural **s**tability *in vivo*.) By labeling known-state structures and evaluating a sequence using these six features, a prediction model can be generated. DIS was used to evaluate the degree of RNA structural destabilization *in vivo*, and its distribution was similar to a normal distribution. To ensure a sufficient number of training sets and development sets (Train/Dev set), we decided to use μ±σ as the threshold value. Class 1 was defined as DIS greater than μ+σ, meaning that such structures had a tendency to become more unstable *in vivo*; class 0 was tagged as DIS greater than μ−σ, indicating that such structures would tend to become more stable *in vivo*. Therefore, the question became a binary classification problem. In this project, we used TensorFlow[28], a deep learning framework created by the Google Brain team, to construct a DNN model to solve the complex problem of RNA structural state prediction *in vivo*. This framework can greatly simplify the process of building the DNN sequential model by using built-in functions. After adjusting the hyperparameters of the model, the final model has 11 hidden layers (10×64 and 1×1), the activation functions adopted were ReLU[29] and Sigmoid, the optimization function Adam[30] was adopted to accelerate the training process, and batch normalization[31] was added to prevent overfitting of the model to the training set (Fig 7a).

**Figure 7.**
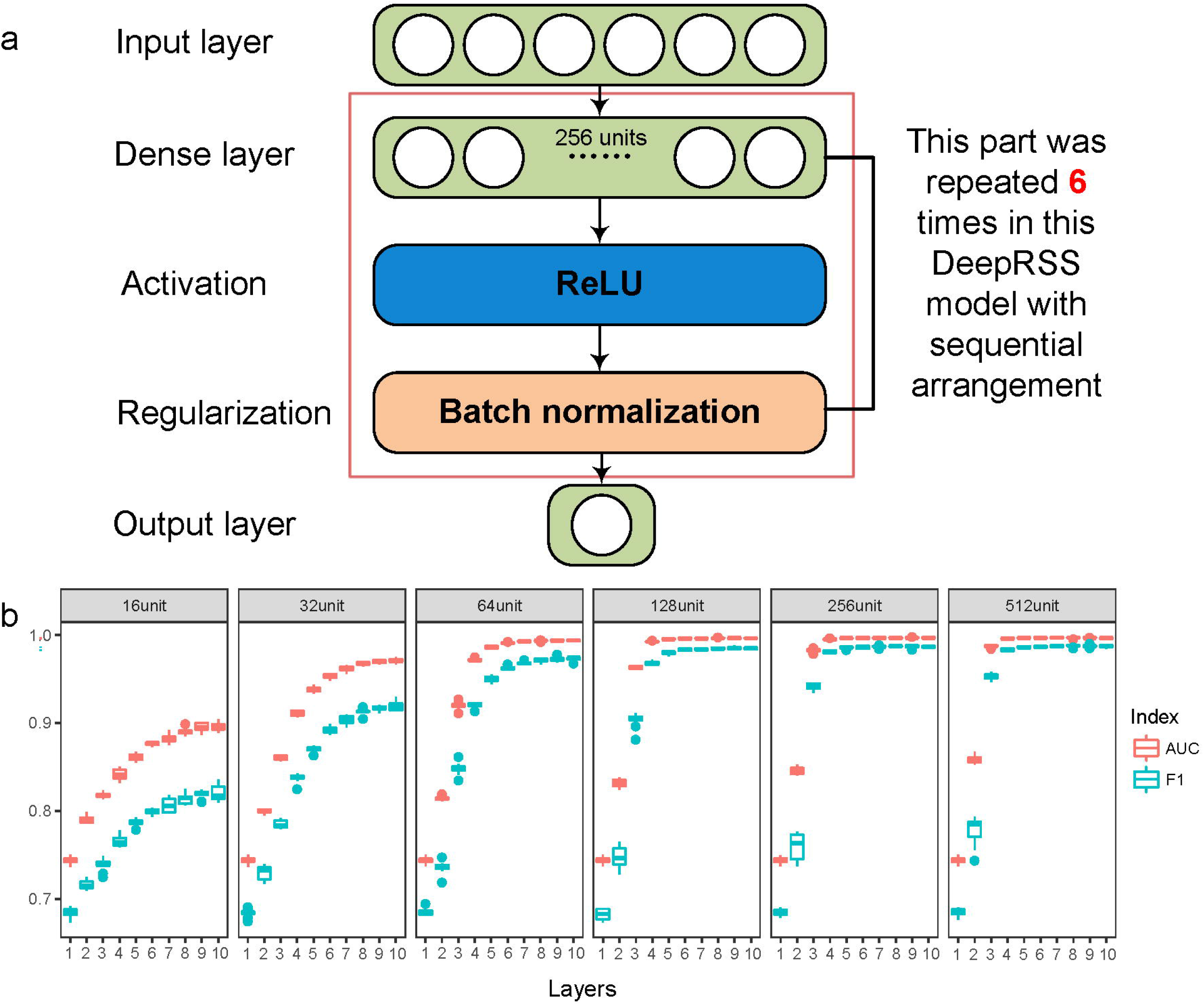
Schematic illustration of the DNN model. (a) The DNN model used in this study is generally a sequential model of multiple dense layers. After careful verification of the number of dense layers and units per layer, we used a sequential model of 6 dense layers of 128 units. More details can be found in the description of the method. (b) Tenfold cross-validation was performed on models constructed with different numbers of layers and different numbers of units.

New paragraph: use this style when you need to begin a new paragraph.

### 10-fold cross-validation

A 10-fold cross-validation (10CV) method was used to evaluate the generalization capabilities of different DeepRSS models. All mRNA structural data (from 135844 mRNA structures) were first randomly shuffled three times (Shell command: cat data | shuf | shuf | shuf >shuffledData) and then randomly divided into 10 groups, each with 9 groups as the training set (Train) and 1 group as the development set (Dev), i.e., this process was repeated 10 times until each group was used once in the development set once. Each model was subsequently trained on a training set for 500 epochs and validated on the development set. To select the most suitable model, we performed a 10CV on models with multiple dense layers (1 to 10 layers) and multiple units per layer (16, 32, 64, 128, 256 and 512 units) (Fig S2-5). The area under ROC curve (AUC) and F1 score (F1) of the 10 cross-validations were averaged.

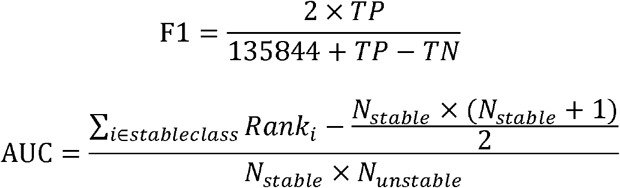

*TP* and *TN* are the true positive and true negative, respectively, of the unstable structure with a threshold of 0.5. *N* is the number of stable and unstable structures. *Rank* represents the index after sorting the data according to the predicted value. The indices of the structures with the same predicted value were averaged to generate the Rank value. The model with 6 fully connected layers of 256 units was chosen as the DeepRSS model because it had the highest AUC value and the lowest SD of the 10 test sets (Fig 7b, Table S1). The training data, Python and Perl scripts, and model files used in this project can be obtained in the Github repository (https://github.com/atlasbioinfo/DeepRSS). Among them, the Python scripts are mainly used for the deep learning model training and prediction, and the Perl scripts are used for the data processing and preliminary analysis.

### Single-factor mutations and predictions

We performed single-factor gradient mutation experiments on the six features of mRNA structure to test the effects of these features on the structural stability. A gradient of 100 aliquots was designed between the maximum and minimum values of the structural features, and gradient mutations were performed from small to large. It should be noted that the significance of the simulated mutation *in silico* was to more accurately analyze the effects of multiple factor effects on the retention of mRNA structures *in vivo*, which were learned by the DeepRSS. First, the *in silico* mutation can design a large number of previously unobservable mRNA structures and predict their characteristics, enriching the amount of data. Second, the value of the turning point of the structural feature’s influence could be obtained by prediction, which is not possible by traditional statistical analyses. When the mutated structure was predicted again by the DeepRSS model, it was known whether the mutation affects the structural state. The transformation of mRNA structure was defined as a change in mRNA structure state (i.e., reclassification into another group) after the mutation. The turning point is the number of mRNA structural state transitions that have occurred in these 100 mutations. For example, a structure was classified into class 0 (stable structure) in the first 40 mutations, and the 41st to 100th were classified into class 1 (unstable structure), which was said to have 1 turning point with the turning point located in the 41st mutation. Because the DeepRSS may increase the number of turning points in the prediction due to the error rate (0.26%), we smoothed out the prediction results of less than 3 consecutive. In the classification based on turning points, the mRNA structures with no turning points (that is, the mRNA structural state did not change during the gradient mutation, named continued stable or unstable structures) could prominently reflect the influence of a certain structural feature on the structural stability state. We judged that the structural features of more than 1 SD of the mean explained the formation of the no turning point mRNA structures. In addition to the POS structural feature, less than 0.2 and greater than 0.8 were chosen as the threshold.

### Impact ratio of the structural features

An mRNA structure in which no structural state transitions (no turning point structures or continued stable or unstable structures) occurred in the gradient mutation of the structural features was selected and classified according to the structural features.

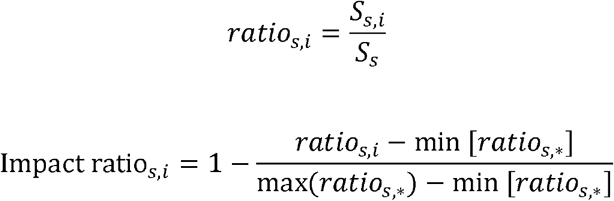

Ratio is the number of continued stable or unstable structures of structural state “*s*” for structural feature “*i*” in addition to the structural state “*s*” total structure “*S*_*S*_”. The impact ratio is the normalized result of all the ratios of a structural state “*s*”.

### Structural feature classification

The 1x and 2x SD are used as the threshold for classification. “exHigh/exLow” refers to structures more than two SD from the mean. “High/Low” refers to structures between “one and two SDs” and “Middle” to “±1 SD structures”. Structural conservation data were obtained from Rouskin’s study[4]. A structural region was randomly extracted from the total mRNA structure, excluding the list of conserved structures. Three parameters that can fluctuate within the cell, RD, RPKM and INI, were varied by a one-fold standard deviation mutation (Up was +SD and down was −SD).

## Acknowledgements

The authors would like to thank the State Key Laboratory of Crop Stress Biology in Arid Areas for its technical and hardware support for this project. We are grateful to Dr. Wenlong Ma for his guidance in deep learning. We thank Dr. Anthony M. Mustoe for advice on mRNA structure analysis and professor Zhao Xu for guidance on data pre-processing. Finally, we thank professor Yanling Liu and Gehong Wei for valuable guidance of the writing of this thesis and critically reviewing the manuscript.

